# *In silico* drug discovery strategies identified ADMET properties of decoquinate RMB041 and its potential drug targets against *Mycobacterium Tuberculosis*

**DOI:** 10.1101/2021.11.17.469062

**Authors:** Kirsten E. Knoll, Mietha M. van der Walta, Du Toit Loots

**Affiliations:** Human Metabolomics, North-West University, Private Bag x6001, Box 269, Potchefstroom, 2531, South Africa

## Abstract

The highly adaptive cellular response of *Mycobacterium tuberculosis* to various antibiotics and the high costs for clinical trials, hampers the development of novel antimicrobial agents with improved efficacy and safety. Subsequently, *in silico* drug screening methods are more commonly being used for the discovery and development of drugs, and have been proven useful for predicting the pharmacokinetics, toxicities, and targets, of prospective new antimicrobial agents. In this investigation we used a reversed target fishing approach to determine potential hit targets and their possible interactions between *M. tuberculosis* and decoquinate RMB041, a propitious new antituberculosis compound. Two of the thirteen identified targets, Cyp130 and BlaI, were strongly proposed as optimal drug-targets for dormant *M. tuberculosis*, of which the first showed the highest comparative binding affinity to decoquinate RMB041. The metabolic pathways associated to the selected target proteins were compared to previously published molecular mechanisms of decoquinate RMB041 against *M. tuberculosis*, whereby we confirmed disrupted metabolism of proteins, cell wall components, and DNA. We also described the steps within these pathways that are inhibited and elaborated on decoquinate RMB041’s activity against dormant *M. tuberculosis*. This compound has previously showed promising *in vitro* safety and good oral bioavailability, which were both supported by this *in silico* study. The pharmacokinetic properties and toxicity of this compound were predicted and investigated using the online tools pkCSM and SwissADME, and Discovery Studio software, which furthermore supports previous safety and bioavailability characteristics of decoquinate RMB041 for use as an antimycobacterial medication.

## Introduction

Tuberculosis (TB), caused by *Mycobacterium tuberculosis*, remains one of the leading causes of death by a single infectious agent (1). Despite the many efforts to search for novel anti-TB treatment, only three drugs have been approved and released into the pharmaceutical market over the last 50 years (2). The inefficiency of many of the existing anti-TB medication, can be attributed to the rapid development of drug resistance and poor understanding of the preceding cellular transition to dormancy (3). Research on compounds effective against active *M. tuberculosis* have shed light on the importance of only a few genes/proteins required for its non-replicative survival, enabling mycobacteria to redirect energy sources for overcoming stress (deprivation of nutrients, acidic environments, and exposure to reactive oxygen species, and various Anti-TB drugs) that resulted in the transition to dormancy and the subsequent anti-TB drug resistance (4). The increasing worldwide prevalence of multidrug-resistant (MDR) and extensively drug-resistant (XDR) TB cases has also highlighted *M. tuberculosis*’s phenotypic plasticity during latency (5–7). Furthermore, drugs that are effective against non-replicating *M. tuberculosis* are still lacking. Additional factors preventing the development of successful anti-TB drugs are the long duration and high costs of novel drug discovery and as the onset of unexpected adverse reactions during clinical trials.

New anti-TB drug discovery is complexed and requires a comprehensive understanding of a drug’s functionality and its target proteins in both *M. tuberculosis* and the host. Fortunately, the ongoing progress of high-throughput laboratory and computational technology has served to promote exponential growth in both data volume and the variety of data sources, which has enabled researchers to create and apply computational prediction techniques, aimed at reducing the lengthiness and cost of all phases of drug discovery (8). Computer-aided drug design entails webservers and software that identify hit targets, analyze target-protein interactions, annotate cellular functionalities of proteins, and anticipate possible side-effects. All have proven helpful during drug development of potent anti-TB compounds or the prevention of the inclusion of toxic drugs in clinical trials (9, 10).

Decoquinate RMB041 (see structure in Figure 1) is an inexpensive compound with promising *in vitro* antimycobacterial activity (11, 12), and some good *in vitro* ADMET (absorption, distribution, metabolism, elimination, and toxicity) properties (13). To identify and characterize decoquinate RMB041’s hit targets, we combined several functions offered by a variety of software and online webservers; reverse pharmacophore docking, also known as target fishing, by PharmMapper (14), prioritization of targets by TDR Targets, identification of correlated proteins by TBDB, mapping of co-expressed proteins by STRING, annotation of metabolomic pathways by KEGG, virtual screening with Discovery Studio (DS) Visualizer, and docking with AutoDock Vina. Virtual docking has presented some difficulties of late, during the analysis of best binding poses (defined as binding affinities) for lead optimization, which is not surprising, considering that there are many factors that must be considered that may influence the binding of molecules (15). Hence, for the purpose of this study, estimated binding affinities were determined (16–18), also considering entropic and enthalpic influences, the mobility of ligands and proteins, the charge distribution over a ligand, surrounding water molecules, and the various possible conformations of the compound of interest (19).

**Figure 1.**
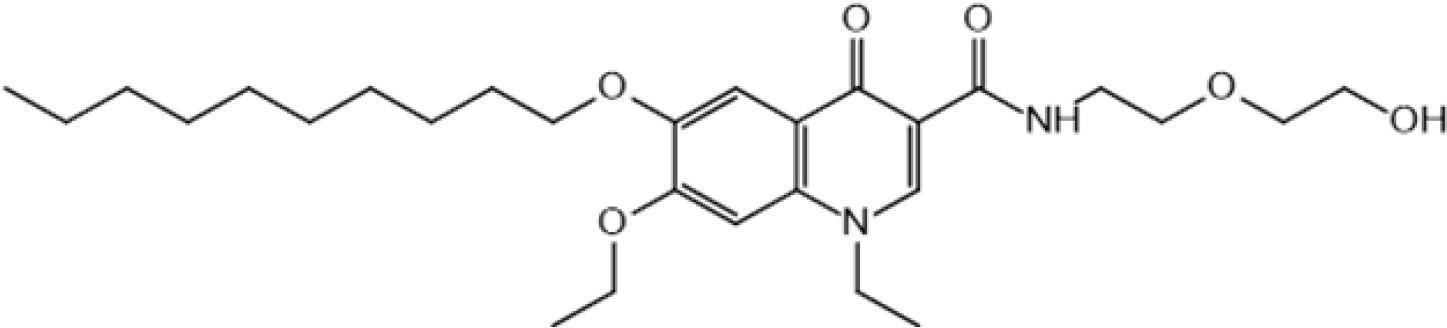
Structure of decoquinate RMB041.

Figure 1. Structure of decoquinate RMB041.

Previously, Beteck, Seldon (12), proved low cytotoxicity of decoquinate RMB041 against WI-38 human fetal lung fibroblasts. In the light of this, we predicted the toxicity of decoquinate RMB041, by investigating those properties generally known to significantly induce toxicity of well-known drugs currently available. Additionally, further properties attributing to absorption, distribution, metabolism, and elimination were determined and evaluated using DS Tools, SwissADME, and pkCSM (20). A previous metabolomic GCxGC-TOFMS investigation of decoquinate RMB041 on *M. tuberculosis*, indicated a mechanism of action by perturbation of the mycobacterial cell wall and DNA synthesis, and also a proposed mechanism by which *M. tuberculosis* might develop a resistance to decoquinate RMB041. In this study, we further contribute to this knowledge by using an *in silico* approach to confirm and build on previous knowledge related to decoquinate RMB041 mechanism of action, toxicity and possible *M. tuberculosis* resistance.

## Results and Discussion

### General molecular properties

Aside from the number of hydrogen acceptors and the number of atoms – the latter because pkCSM counts only heavy atoms, while DS counts all atoms of the molecule – SWISSadme and DS agreed on all decoquinate RMB041’s molecular properties (Table 1).

**Table 1.**
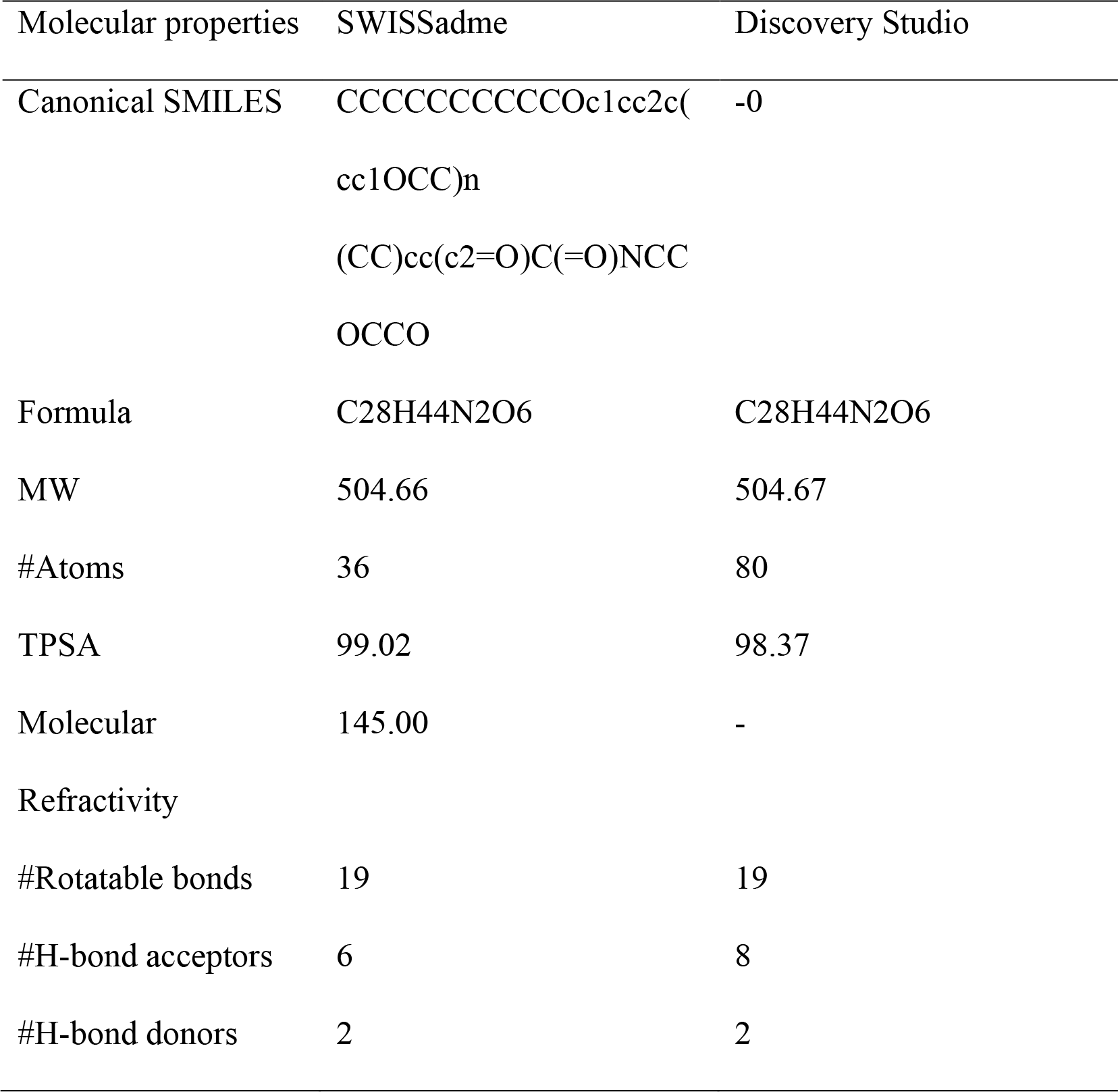
Molecular properties of decoquinate RMB041, as provided by SWISSadme and Discovery Studio.

### Likeness of oral administration

Pharmaceutical companies generally apply one or more regulations that assist in determining the drug likeness of oral administration, of which Lipinski’s rule of five is the most common criteria (21). The number of violations of the rules of Lipinski, Veber, Egan, and Muegge, provided by SWISSadme, along with the molecular properties breaching these rules, are presented in Table 2. With regards to the applied pharmaceutical regulations, only the molecular weight (MW) and the amount of rotatable bonds breach the rules. These broken rules do not necessarily indicate a lack of drug efficacy (22). All in all, these results indicate promising effectivity of decoquinate RMB041 after oral administration.

**Table 2.** Number of violations of commonly applied pharmaceutical rules of drugability, as provided by SWISSadme.

| Pharmaceutical test | SWISSadme | Rule violated | Reference |
| --- | --- | --- | --- |
| Lipinski #violations | 1/5 | MW > 500 | Lipinski<br>(23) |
| Veber #violations | 1/2 | #Rotatable bond > 10 | Veber,<br>Johnson<br>(24) |
| Egan #violations | 0/2 | - | Egan,<br>Merz (25) |
| Muegge #violations | 2/8 | MW > 300<br>#Rotatable bond > 15 | Muegge,<br>Heald (26) |

**Table 3.**
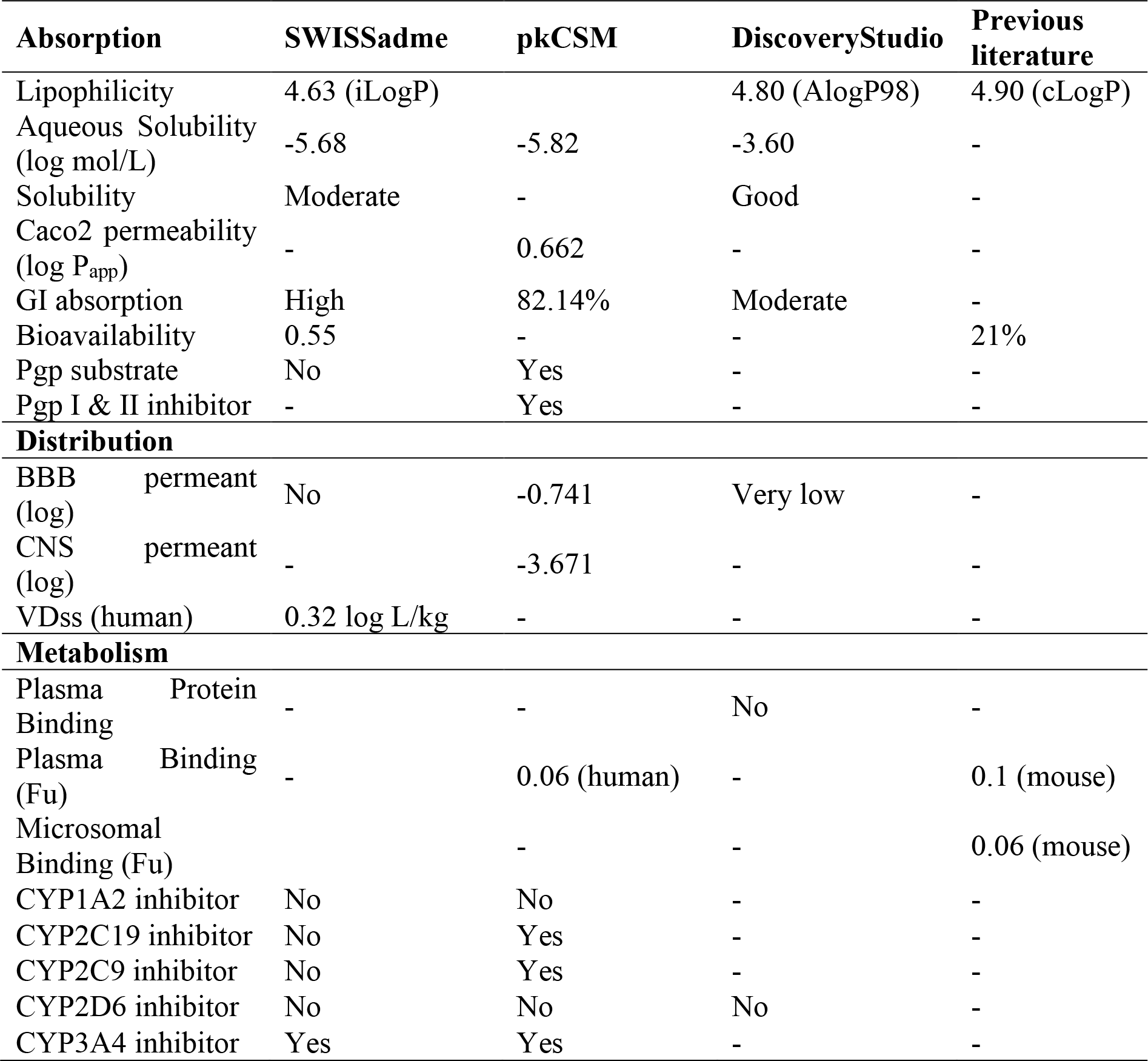
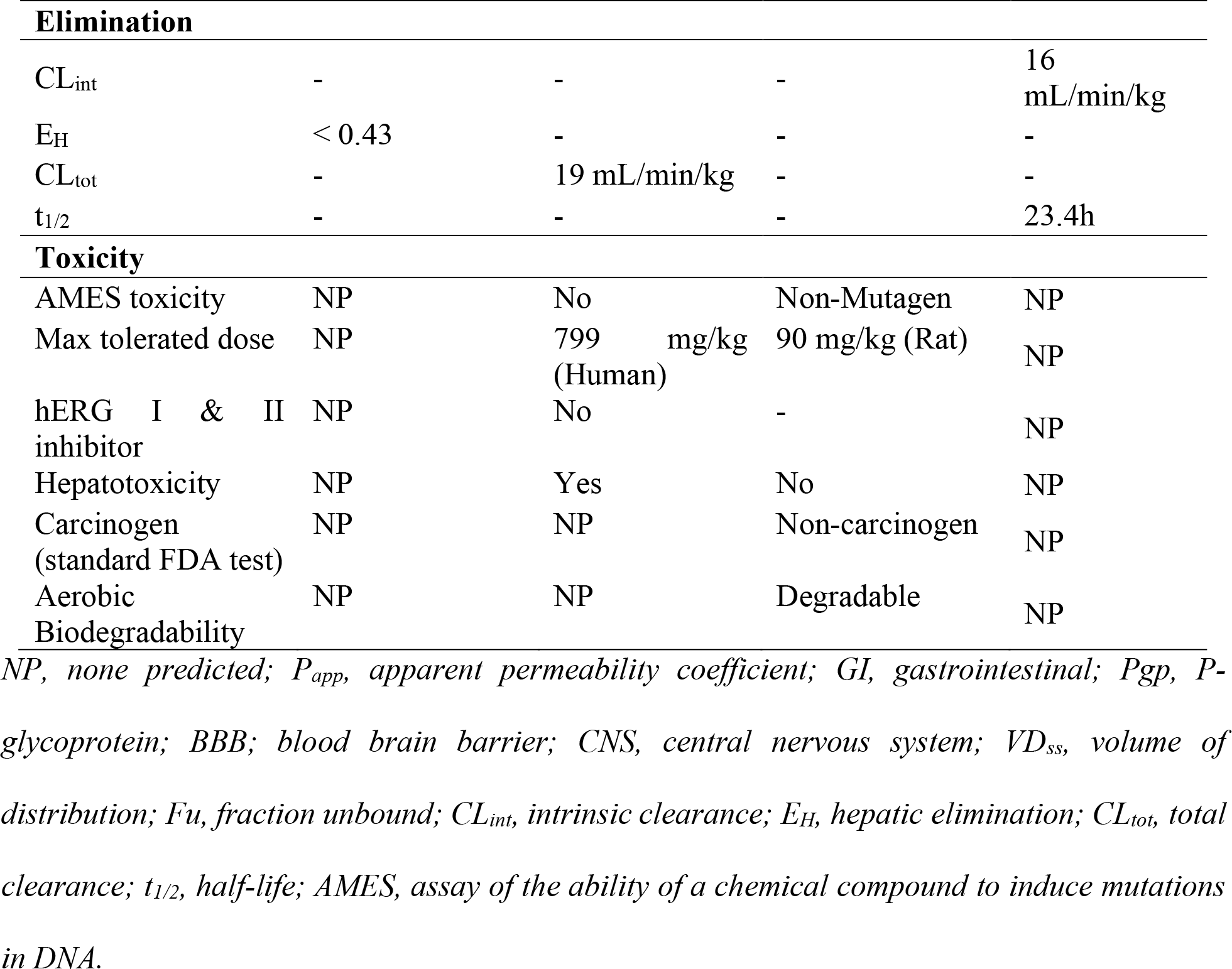
ADMET pharmacokinetic properties of decoquinate RMB041, provided by computational prediction methods and previous literature.

**Table 4.**
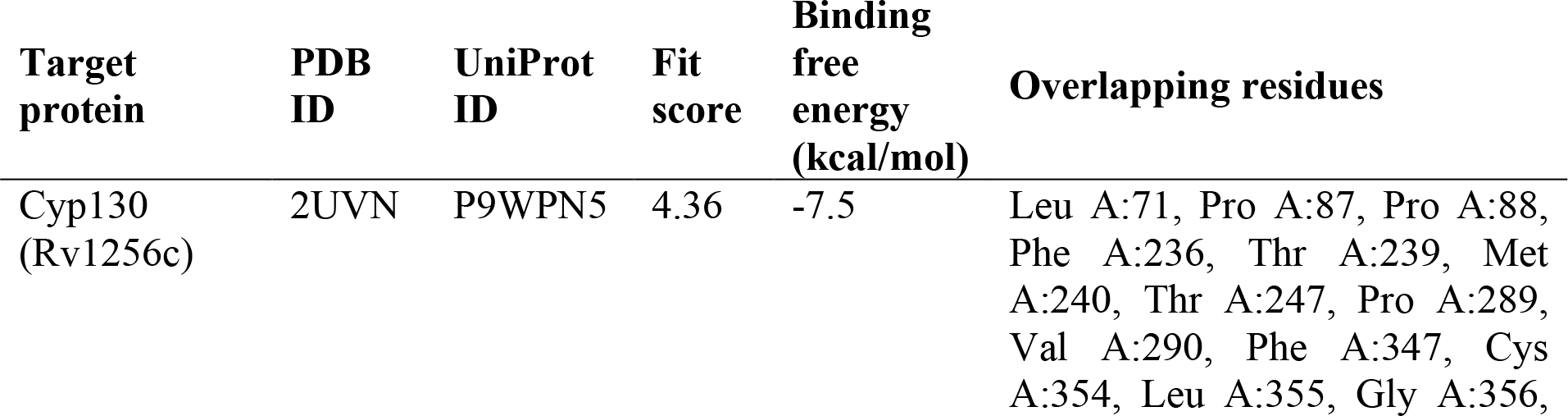
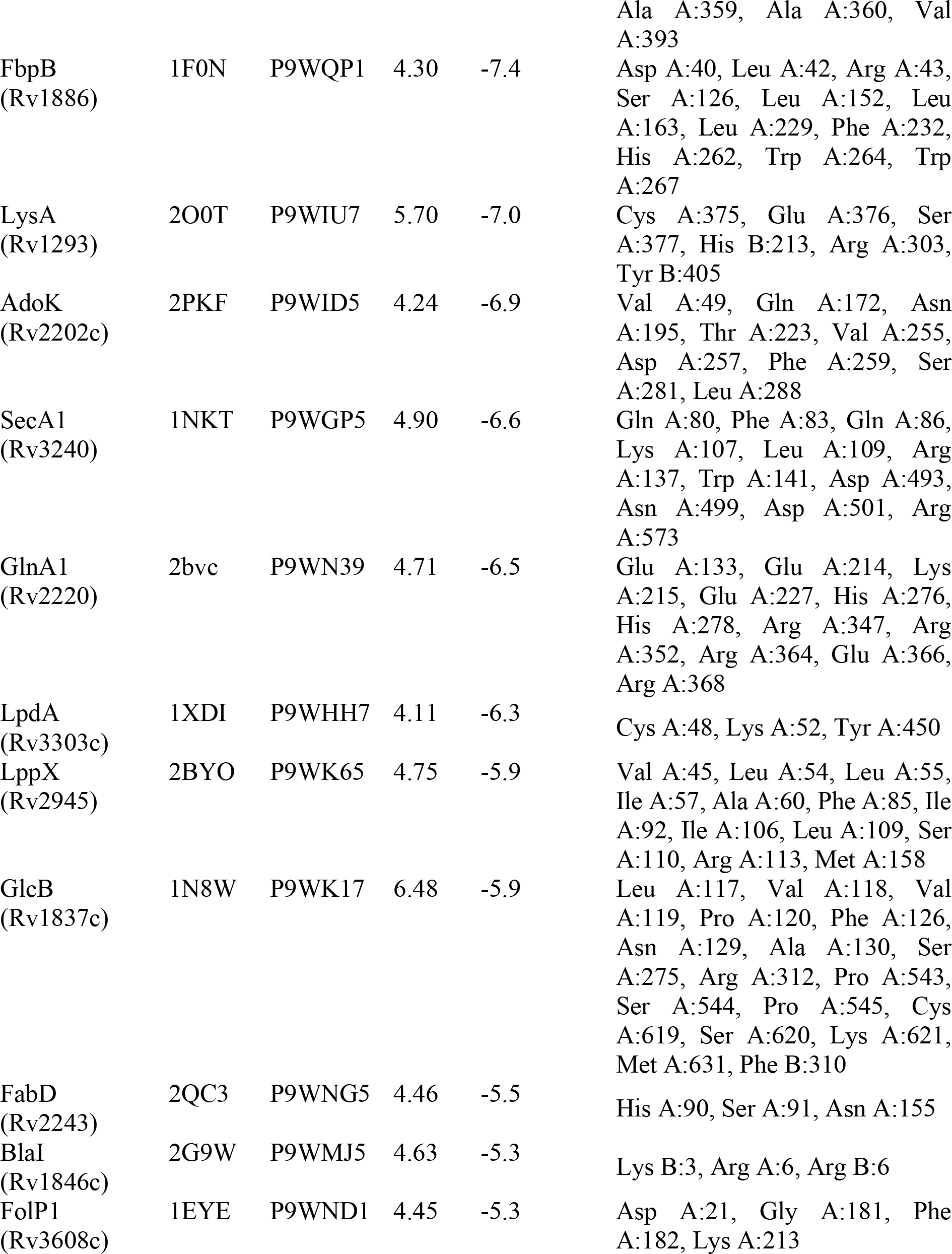
The targets identified by PharmMapper, along with their respective identification codes, fit scores, binding free energies, and residues that interact with both, their respective co-crystalized ligands and decoquinate RMB041.

### Absorption, distribution, metabolism, elimination, and toxicity

In earlier studies, decoquinate RMB041 (MIC_90_ = 1.25 μM) has shown to have promising bioavailability and distribution characteristics (13). Additional positive ADMET-related properties were also indicated by DS and prediction webservers used in this study. ADMET influencing factors from these tools and previous literature (12, 13) are presented in **Error! Reference source not found.**.

Considering all parameter related to decoquinate RMB041 absorption, the high lipophilicity (LogP = 4.63 to 4.90; LogD = 4.80) of decoquinate RMB041 suggests excellent permeation through the mycobacterial cell wall and into macrophages, and likeliness of being more effective against dormant *M. tuberculosis* (27). Moderate to good aqueous solubility (**Error! Reference source not found.**) indicates its practicality during drug formulation and gastrointestinal (GI) absorption (28, 29). A bioavailability score of 0.55 additionally confirms good absorption after oral administration (Martin YC 2005). However, the low Caco-2 cell permeability (human model for GI absorption; High when LogP_app_> 0.9) indicates that further improvement of the solubility might be necessary. Another important influencer on absorbance is the amount of efflux from cellular tissue, especially by p-glycoprotein (Pgp). The contradictory results (SWISSadme vs pkCSM) regarding the binding of decoquinate RMB041 to Pgp (**Error! Reference source not found.**) suggests further experimentation regarding this would be necessary. In either case however, whether it is not a substrate for Pgp (as predicted by pkCSM), or whether it is a substrate, and inhibits Pgp I and II, the results indicate a lack of export (which parameter would this be?), meaning decoquinate RMB041 is likely to accumulate in the target organs.

When considering the distribution criteria, all three prediction methods state unlikely penetration of decoquinate RMB041 through the blood-brain barrier (BBB) (**Error! Reference source not found.**) (30). This may indicate a low probability of neurotoxic effects, such as confusion, depression, psychosis, or muscular weakness, which appear in response to various other current anti-TB medications (31, 32). It may also indicate however, that this compound would likely be of little use for the treatment of TB meningitis. Blood plasma volume of distribution of decoquinate RMB041 (log VDss > 0.32) (**Error! Reference source not found.**) is on the higher end of the spectrum (log VDss < -0.15 is low; log VDss >0.45 is high), suggesting moderate delivery to infected areas (33, 34).

Considering the metabolism of the decoquinate RMB041, the cytochrome P450’s, the most significant metabolizing enzymes to be considered during drug metabolism, oxidize xenobiotics to facilitate their excretion (35). The inhibition of these enzymes by an antibiotic indicates the likelihood of toxic accumulation, as well as possible interference with the pharmacokinetics of co-administered antibiotics. Despite differing conclusions about CYP2C9 and CYP2C19 inhibition by decoquinate RMB041, pkCSM’s and SWISSadme’s calculations regarding the two main cytochrome isoforms, 2D6 and 3A4, are in agreement (**Error! Reference source not found.**). CYP2C9 was recognized by PharmMapper as a potential protein target, supporting the results by pkCSM. Although no inhibition of CYP2D6 is expected, inhibition of CYP3A4 suggests that care should be taken when co-administering decoquinate RMB041 with other antimycobacterial compounds, in order to prevent toxic accumulation of either or both drugs.

When evaluating the elimination characteristics of decoquinate RMB041 using the data generated by pkCSM, SWISSadme, and previous experimentation results by Tanner, Haynes (13) and colleagues, we retrieved estimated values for the total clearance rate (CL_tot_ = 19 mL/min/kg), intrinsic clearance (reflecting hepatic and biliary excretion) (CL_int_ = 16 mL/min/kg), hepatic excretion (E_H_ < 0.43), and the elimination half-time (t_1/2_ = 23.4 hours) (**Error! Reference source not found.**). The t_1/2_ is far superior to those of the other first-line drugs rifampicin (t_1/2_ = 7 hours) (36), isoniazid (t_1/2_ = 1.7 hours) (37), and ethambutol (t_1/2_ = 3 hours) (38), and indicates less frequent drug administration would be required, which is beneficial for patient coherence and possibly decreasing the consequential occurrence of drug resistance.

Decoquinate RMB041 has previously shown little cytotoxicity against WI-38 human fetal lung fibroblasts (IC_50_ = 56.2 μM) (12). According to pkCSM and DS, this compound is neither carcinogenic, nor mutagenic, nor likely to interfere with heart rhythms (**Error! Reference source not found.**). Furthermore, the odds of having neurotoxic and hepatotoxic properties are also low. The positive hepatotoxicity by pkCSM is only based on the similarity of structural features to compounds with liver-associated adverse effects, whereas DS, which showed no possible hepatotoxicity, uses additional information related to dose concentrations to establish the probability of hepatotoxicity. According to the info generated in this investigation, Decoquinate RMB041 shows a promising safety profile compared to that of other first- and second line antitubercular drugs which exibit cardiotoxic, hepatotoxic and/or neurotoxic effects (32, 39–41). That said, the contradictory results by pkCSM and DS concerning hepatotoxicity necessitate further experimentation. Aerobic biodegradability, a trait that is often underestimated and missed during drug development, is important for the prevention of wastewater pollution, which would else cause serious harm to the aquatic ecosystems and increase antibiotic resistance in humans (42). The degradation of this compound therefore suggests that it is safe in the face of environmental pollution also, just as a matter of interest.

### Identification and prioritization of drug targets

It is widely known that drugs commonly target several proteins (as opposed to only one particular protein) (43). PharmMapper compares the pharmacophores of the investigated drug compound, to those derived from ligands in complex crystal-structures and provides predicted corresponding protein targets (14). A total of twelve *M. tuberculosis* proteins were identified by PharmMapper as potential targets of decoquinate RMB041. These are listed according to their docking scores in Table , along with their respective PharmMapper fit scores, UniProt and PDB accession codes, and associated KEGG reactions. A limitation one should keep in mind about current computer-aided drug design, is that only the known active sites in databases can be used for identifying suitable ligand scaffolds. Further research on active sites of proteins would increase the accuracy of the elucidated mechanism of action, which will undoubtable improve as more data becomes available regarding this.

The docking affinities calculated by AutoDock Vina ranged between -7.5 to -5.3 kcal/mol and indicated strong binding to the target proteins (Table ). Several studies have ranked targets by their importance in *M. tuberculosis*, focusing mainly on drugability and essentiality *in vitro* survival (44, 45). However, it is also important to consider the various conditions that *M. tuberculosis* are exposed to in the host during infection or disease, such as acidity, reactive oxygen species, nutrient restriction, and various antibiotics in the case of a treated TB patient. During genomic, proteomic, and metabolomic investigations of *M. tuberculosis* exposed to each of the aforementioned conditions, it has become apparent that, although the initial response mechanism may differ, *M. tuberculosis* survival ultimately depends on its ability to persist in a non-replicative/latent TB state, which is the case in most individuals infected with TB, which is for one third of the global population currently (1), and hence, extremely valuable to investigate targets that are essential for *M. tuberculosis* persistence. One should also consider the neighboring network of the target gene/protein, i.e., targeting a gene/protein associated with many other gene/proteins might prove more successful than targeting one with fewer interacting genes/proteins. Lastly, the exclusivity of a target gene/protein to *M. tuberculosis,* is also an extremely sought-after characteristic in drug design, and additionally, if other genes/proteins within in *M. tuberculosis* have a similar function, its less likely to disarray essential cellular processes.

In order to predict the decoquinate RMB041’s drug targets, we determined their ranking by TDR Targets, which, with the help of integrated databases, allows users to prioritize genes and their annotated proteins based on various filtering criteria and criteria-specific weighting (46). We filtered non-human homologs and looked at three ranking orders: (1) with respect to homology to gut flora, similarity to essential genes, and maintaining of persistence (47), (2) upregulation of genes during dormancy and transition to dormancy (48), and (3) ortholog-based inference of essentiality of genes during experimental conditions relevant to drug discovery. Based on the first ranking order, 313 protein targets annotated to highly prioritized genes were provided, of which only one, Cyp130, was also identified by PharmMapper as a target of decoquinate RMB041. The second-ranking order provided a list of 420 proteins, of which only one, BlaI, is a decoquinate RMB041 target. According to the third-ranking criteria, both Cyp130 and BlaI (2 out of 397 proteins) are essential targets of decoquinate RMB041. *FolP1* and *lysA* were also number one and number three, respectively, on a list of 4000 genes that were ranked based on their metabolic uniqueness in a study by Hasan, Daugelat (47), further indicating the importance of their encoded proteins.

#### Cellular mechanisms

The KEGG database BRITE hierarchy files annotated ten of the twelve targets, of which all are involved in *M. tuberculosis* energy metabolism, protein synthesis, fatty acid synthesis, cell wall synthesis, TCA cycle activity, and/or amino acid synthesis. Every one of these pathways were found altered in the previous study (11). Correlations of *M. tuberculosis* genes have been extensively studied and are still barely understood. Aside from the very many genes in *M. tuberculos*is (49), the complexity of interdependent intracellular genetic expressions makes it almost impossible to predict the complete cellular response to external stimuli accurately. For instance, even if the expressed proteins show a strong positive correlation to each other, it is impossible to tell if the inhibition of one would lead to the inhibition of the other, without extensive experimentation of each interaction separately. However, it can indicate the metabolic pathways that are likely to be altered in the inhibition process and give information on which drugs might work well in conjunction with decoquinate RMB041. With help of STRING and TBDB, we identified protein targets and analyzed direct correlations between these (Figure 2). TBDB links *M. tuberculosis* gene-expression microarray data to their encoded proteins and provides the strength of the correlation between proteins (50). Of a total of 144 correlated proteins, only ten were negatively correlated. If one were to assume that inhibition of the protein targets would influence the expression of the correlated proteins, it would mean that 134 other proteins might be expressed to a lesser degree, while ten proteins might be expressed to a larger extent.

**Figure 2.**
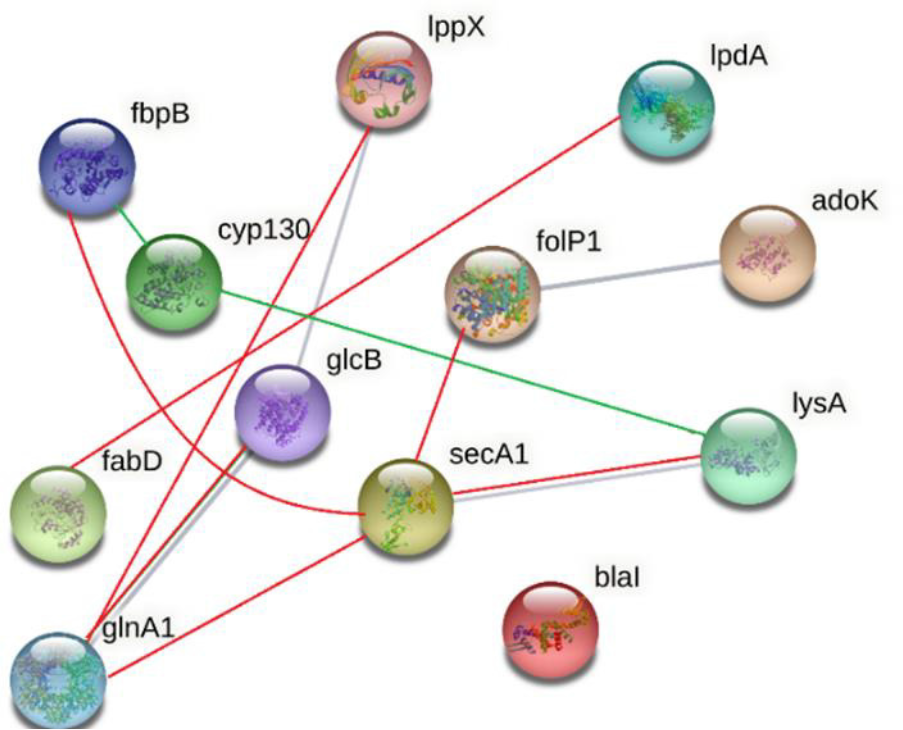
Interconnected network provided by STRING (grey), with additional interactions retrieved from TBDB; Positively correlated interactions (red) and negatively correlated interactions (green).

Beteck, Seldon (13) and colleagues established that decoquinate RMB041 primarily acts on the cell wall and, secondly, inhibits DNA synthesis. Our findings during an earlier metabolomic investigation confirmed this and also suggested that decoquinate RMB041 inhibits protein synthesis, and pathways associated with *M. tuberculosis* in the non-replicating state (11). Here, we examined each drug target to better understand the perturbations caused by decoquinate RMB041 and explore the cellular response by which *M. tuberculosis* counteracts these, such as those leading to the metabolomic changes revealed in our previous investigation.

BlaI controls the expression of the Rv1864c regulon, which comprises genes involved in drug transport, detoxification, and ATP synthesis (51). Dissociation of this enzyme from its operator site allows the transcription of the genes belonging to this regulon, including *blaI,* encoding itself, and *blaC*, encoding beta-lactamase (BlaC) (52), which causes resistance to beta-lactams (53). Depending on the ratio of BlaI inhibition versus *blaI* transcription and the mycobacterial degradation of the inhibitor, it could either enhance efflux, ATP synthesis and tolerance to stress, or repress the regulon, leading to opposite effects. Either way, it would disrupt the NAD(P)H: NAD(P)^+^ ratio. Considering that previous studies indicated elevated NAD(P)^+^ in the presence of decoquinate RMB041 (11), it is tempting to assume that hydrogen ions are lacking, meaning ATP synthase is inhibited, of which one could deduct that the entire regulon is being repressed. Considering that TDR Targets also deducts its answers based on orthologous experimental results of studies on other bacteria, such as *Eschericia coli* or staphylococci, the prioritization of BlaI as a drug target is a promising outcome. Nonetheless, further experimentation would bring clarity to what the consequences of BlaI inhibition would be. The DNA binding site of BlaI shares three residues with that of decoquinate RMB041 (based on the AutoDock Vina results), and the interaction between the drug and BlaI involves five strong hydrogen bonds (Figure 3. Interactions between (A) Cyp130 and decoquinate RMB041, (C) BlaI and decoquinate RMB041, and (D) LpdA and decoquinate RMB041.Figure 3).

**Figure 3.**
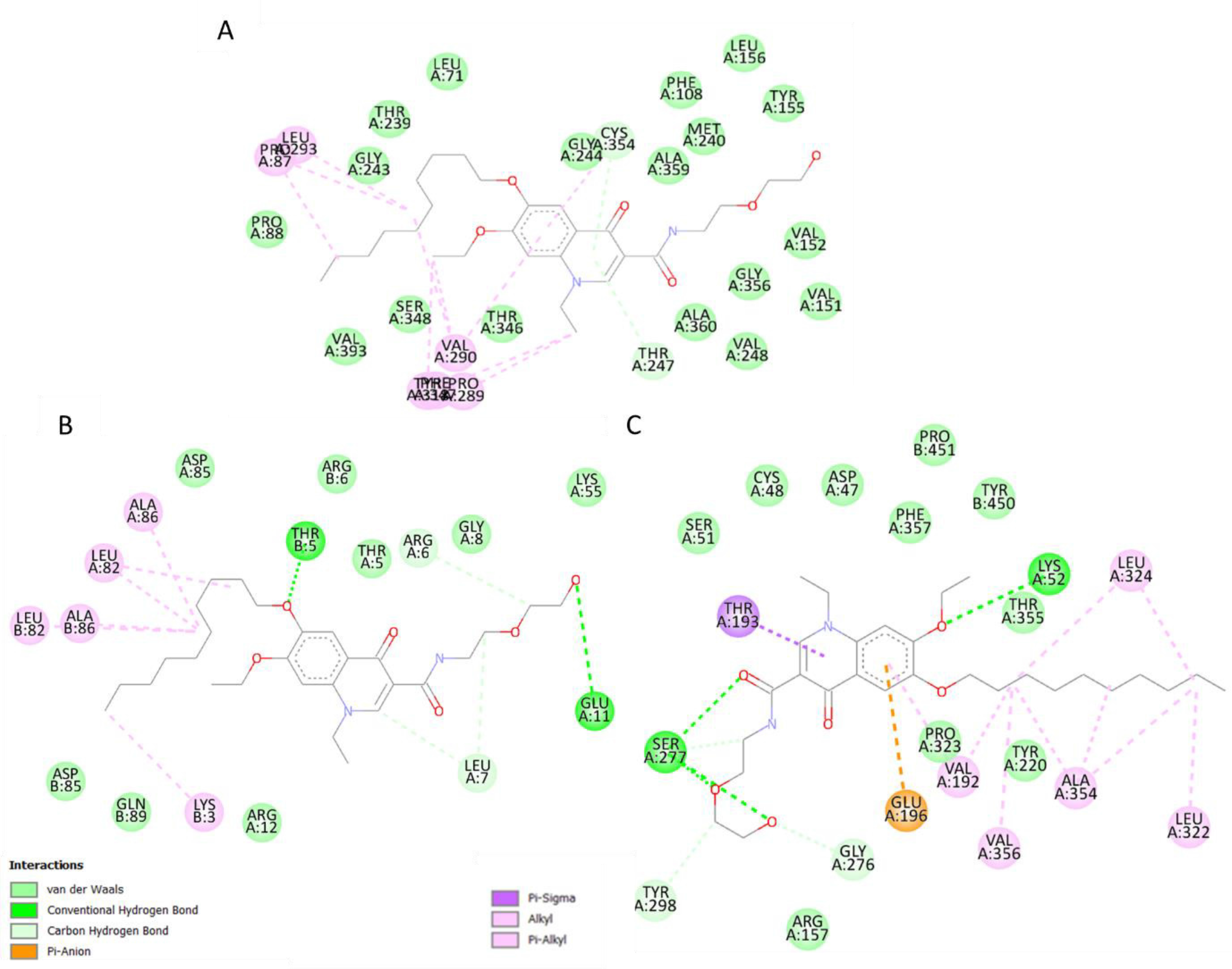
Interactions between (A) Cyp130 and decoquinate RMB041, (C) BlaI and decoquinate RMB041, and (D) LpdA and decoquinate RMB041.

Cyp130 is one of twenty cytochrome P450 enzymes responsible for the incorporation/reduction of molecular oxygen, and functions in conjunction with Tap (Rv1258c), for the export of drug molecules (54). Furthermore, it is well known that Cyp130, and its encoding gene, *cyp130*, are crucial for growth and virulence of *M. tuberculos*is, and the inhibition of Cyp130, effective against MDR-*M. tuberculos*is strains (49). In the aforementioned study, Cyp130 was also indicated with a high priority as an anti-tuberculous target in two of the selected TDR Target ranking orders, and reportedly inhibited by a drug group called azoles, with econazole showing the most potent activity (49). Interestingly, the binding cavity of econazole includes sixteen residues that also interact with decoquinate RMB041 (55) (Figure 3). Cyp130 shows the highest comparative binding affinity to decoquinate RMB041, mostly via hydrophobic interactions and two strong hydrogen bonds.

LpdA is a NAD(P)H-requiring flavoprotein disulfide reductase that has been found to be significantly upregulated during the transition of *M. tuberculos*is towards a state of dormancy (56, 57). Within the active cavity of LpdA, bound to both, FAD and NADP+, three residues also interact with decoquinate RMB041 (Figure 3). Inhibition of this enzyme would prevent the conversion from 2,6-dimethyl-1,4-benzoquinone to 5-hydroxy-1,4-naphraquinone and a subsequent accumulation of NAD(P)H. Elevated NAD(P)H levels in turn, would induce counteractive pathways, for instance the NAD(P)H dependent production of α-hydroxyglutaric acid, fatty acids, and sugar alcohols. All three pathways have been found to be altered in our previous metabolomics study (11), further supporting decoquinate RMB041’s inhibition of LpdA and, accordingly, a promising activity against dormant/latent *M. tuberculosis*.

Two of the identified target proteins, SecA1 and LppX (Figure 4), are transmembrane transporters. SecA1 is ATPase coupled and responsible for the transport of the majority of proteins, including those suppressing phagocyte maturation (58). Its previously demonstrated active site, when bound with adenosine diphosphate (ADP)-β-S, shares eleven residues with the binding site of decoquinate RMB041 (59). Inhibition of SecA1 results in an elevated ATP:ADP ratio, i.e., which would disrupt intracellular energy metabolism, in addition to prevent the export of proteins required for cell wall biosynthesis (60, 61). The assimilation of these proteins would in turn induce protein degradation, especially during nutrient starvation (62), to provide proteinogenic amino acids as a source of nitrogen and carbon. This response is supported in our previous metabolomics study, which indicated elevated levels of proteinogenic amino acids in decoquinate RMB041 treated *M. tuberculosis* (11). SecA1 has been referred to as an optimal co-target, with “co-target” defined as a protein whose inhibition, in addition to that of the primary target, would hinder the development of resistance (63). The other transmembrane transporter, LppX, functions by carrying lipophilic molecules across the mycobacterial membrane. Its encoding gene, *lppX,* is upregulated during host-infection (64) and enhances the bacilli’s capability to escape the host’s immune response (65). Decoquinate RMB041 and vaccenic acid, share nineteen residues within the active cavity site with which they interact (65). Compared to vaccenic acid, decoquinate RMB041 forms several more hydrophobic interactions, indicating a stronger competition. Inhibition of LppX, would likely cause accumulation of fatty acids, confirmed by the drastically elevated levels of these compounds in our previous metabolomics study (66).

**Figure 4.**
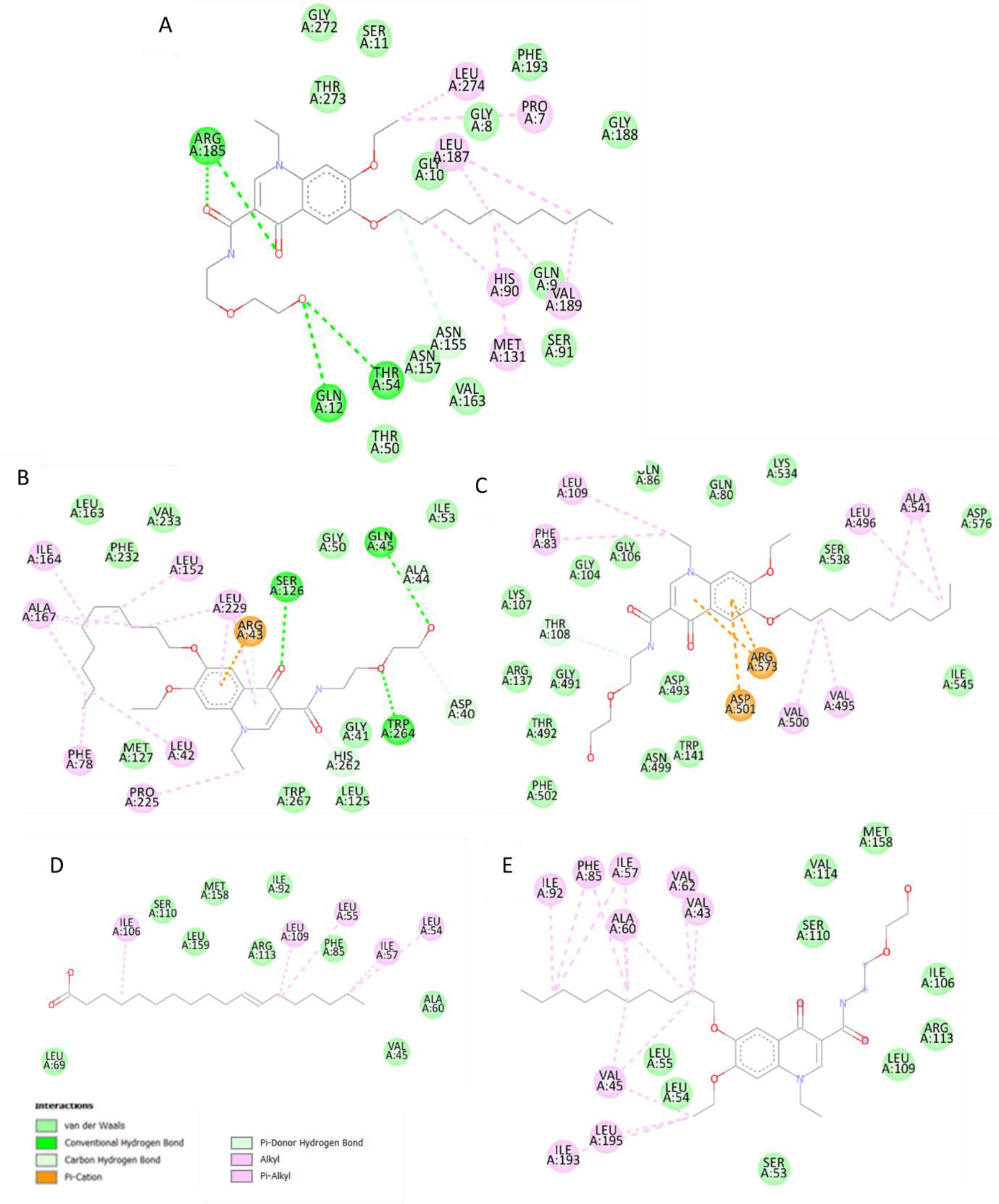
Interactions between decoquinate RMB041 and (A) FabD, (B) FbpB and (C) SecA1, as well as those between LppX and (D) vaccenic acid and (E) decoquinate RMB041.

FabD, the malonyl transacylase, is the first of five enzymes (fabD, acpM, kasA, kasB, and accD6) responsible for fatty acid elongation (67, 68). The role of the encoding gene, fabD, during the stress response is unclear, as it was underrepresented after exposure to rifampicin (69), upregulated in the presence of isoniazid (70), and overexpressed after the removal of a stressor (71). Nonetheless, it has frequently been associated with drug resistance (70, 72), indicating it’s importance for survival after exposure to antibiotics. FabD is a target of the Pup-proteasome degradation (73), which could mean that the protein is readily marked for destruction, followed by the regeneration of mutated FabD. When FabD is inhibited, malonyl-CoA accumulates (12), and also preferentially used as a substrate for mycolic acid synthesis, an important cell wall component for survival during dormancy (74). Despite this however, no shortage of elongated fatty acids occurs, indicating the existence of similarly functional proteins, and/or the break-down of cell wall components into fatty acids, that are considered less vital for survival under conditions of stress, which in this case were trehalose dimycolates. This enzyme’s lack in unique functionality, would also explain why it was not considered as an important target by TDR Targets. The active site of the original crystal structure of FabD contains acetic acid (75), of which three interactive residues also bond with decoquinate RMB041 (Figure 4). FabD is positively correlated to KasA, an enzyme involved in the inhibitory activity of isoniazid on fatty acid elongation, suggesting possible synergistic activity of decoquinate RMB041 with isoniazid.

FbpB, also known as antigen 85B, catalyzes the biogenesis of trehalose dimycolates (TDM) (catalo 19) and is the most abundant of all mycoyl transferases. TDM accumulate outside the cell wall and desensitize the cell to antibiotics (76). FbpB is overexpressed during early infection of macrophages, suggesting it to be important for the shift to *M. tuberculosis* dormancy/latency (77). Inhibition of FbpB would result in the accumulation of TDM, followed by its break-down into mycolic acids, which can either be used to synthesize other cell wall components or be broken down further into long-chain fatty acids. Although FbpB is highly expressed during early infection of macrophages (78), its encoding gene, *fbpB*, has been seen to be downregulated during hypoxia (79), indicating little importance of FbpB during dormancy. Nonetheless, FbpB inhibition might serve useful to prevent desensitization of co-administered antimycobacterial agents. The binding of decoquinate RMB041 to FbpB (Figure 4) is verified by eleven interacting residues that also non-covalently bind to trehalose (78). Interactions between decoquinate RMB041 and FbpB, showing the second highest binding affinity, include six strong hydrogen bonds.

Malate synthase (GlcB) is responsible for the synthesis of malic acid via the glyoxylic shunt, a key pathway during *M. tuberculosis* dormancy, that incorporates even chain fatty acids into the TCA cycle (80). GlcB upregulates the dormancy regulator (DosR) regulon and increases tolerance to stress (81). Inhibition of GlcB would explain the accumulation of even-chain fatty acids shown in our previous metabolomics study on decoquinate RMB041 treated *M. tuberculosis*, although levels of malic acid were increased, likely due to the incorporation of proteinogenic amino acids (11). Although GlcB was noted as as necessary during *M. tuberculosis* dormancy/latency, its absence in the prioritizing lists of TDR Targets indicates that *M. tuberculosis* possesses several enzymes controlling malic acid. The latter could be expected, considering that it is a primary element within the central carbon flux. GlcB’s natural ligand, acetyl coenzyme A, shares seventeen interacting residues with decoquinate RMB041 (Figure 5) (82). According to TBDB, GlcB is positively correlated to rifampicin targets, RpoB and RpoC, suggesting possible synergistic activity of this drug with decoquinate RMB041.

**Figure 5.**
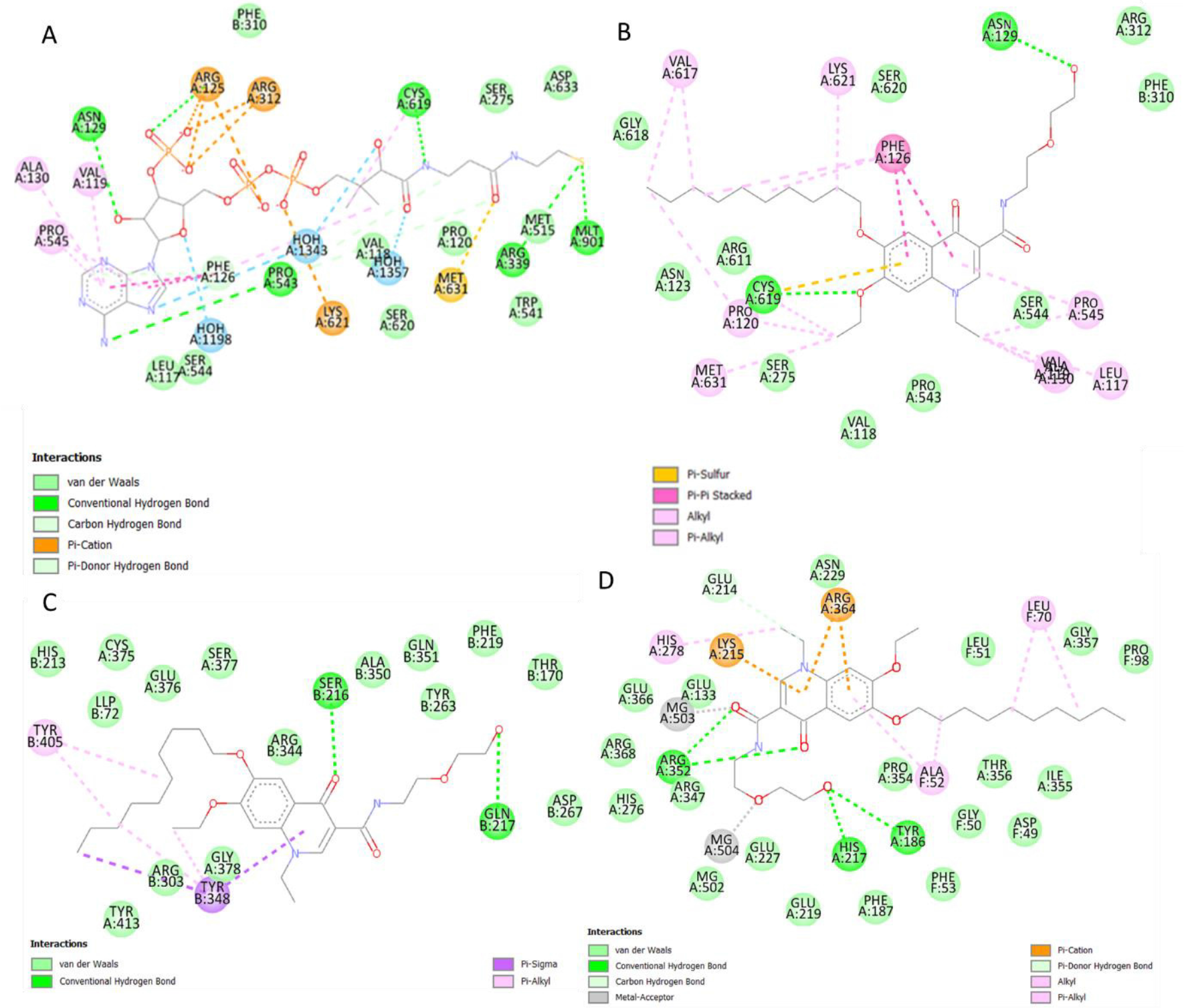
Intercations between (A) GlcB and acetyl-coenzyme-A, (B) GlcB and decoquinate RMB041, (C) LysA and decoquinateRMB041, and (D) GlnA1 and decoquinate RMB041.

*glnaMeso*-diaminopimelate decarboxylase (LysA) catalyzes the final step of lysine biosynthesis. Its encoding gene, *IysA*, is regulated by LysR, which is repressed in the presence of lysine (83). Upregulation of *lysA* has been noted in immunocompromised mice (23), and the highly conserved nature of this gene indicates that it is mainly associated with *M. tuberculosis* stress responses (49). Under normal circumstances, inhibition of LysA would lower lysine levels and LysR would keep inducing lysA expression, creating a metabolic loop and preventing protein synthesis. However, concurrent protein degradation would provide sufficient lysine for the organism, as shown in our previous metabolomics study, which would in turn repress LysR. Hence, inhibition of LysA would, in this case, not provide much assistance in eliminating mycobacteria. The targeting of this enzyme is supported by six residues that interact with both decoquinate RMB041 and lysine (Figure 5) (84).

Glutamine synthetase (GlnA1) is a key enzyme used during nitrogen metabolism and cell wall synthesis (85). It catalyzes ATP-dependent ammonium condensation with glutamate to form glutamine, adenosine diphosphate, and phosphate. The crystal structure of GlnA1 was investigated in a complex with methionine sulfoximine phosphate, a product of methionine sulfoximine phosphorylation, and includes eleven residues in its active binding site that also appear in the binding pocket of decoquinate RMB041 (Figure 5)(86). Inhibition of this enzyme would explain the previously identified elevation of glutamate levels (11). GlnA1 has been proposed as a promising anti-TB target with high importance during dormancy (76, 87) and mycobacterial growth in macrophages (88). According to TBDB, GlnA1 is positively correlated to four proteins that are targeted by other anti-TB drugs. RpsL, EmbA, RpsA, and AtpE are drug targets of streptomycin (protein synthesis inhibitor), ethambutol (mycolic acid transfer inhibitor), pyrazinamide (fatty acid synthesis inhibitor), and bedaquiline (ATP synthase inhibitor), respectively. Hence, further experimentation of decoquinate RMB041 with one or more of these drugs might prove synergistic activity against *M. tuberculosis*.

Adenosine kinase (AdoK) catalyzes the phosphorylation of adenosine to adenosine monophosphate (AMP). This enzyme participates in the purine salvage pathway and plays a crucial role in nucleotide synthesis during *M. tuberculosis* persistence (89). The active binding site was previously illustrated with adenosine (90) and involves nine residues that also interact with decoquinate RMB041 (**Error! Reference source not found.**). Inhibition of this enzyme would be expected to prevent the formation of DNA during the shift to dormancy (as opposed to acutely disrupt DNA synthesis when administered). This “belated” disruption of DNA formation by decoquinate RMB0041, was also confirmed in two previous studies (11, 12).

Inhibition of folic acid synthase (FolP1) would also lead to a disrupted DNA synthesis in *M. tuberculosis*, as proven by previous experiments on mycobacteria in the presence of sulfamethoxazole and trimethoprim (91). FolP1 catalyzes the synthesis of the 6-hydroxymethyl-dihydropteroate and pyrophosphate, from the substrates 6-hydroxymethyl-7,8-dihydropterin-pyrophosphate and *para*-aminobenzoic acid and has previously been proposed to be a promising drug target against *M. tuberculosis* (91). Its encoding gene, *folP1*, is upregulated during the early and late dormancy (56), indicating its importance in maintaining *M. tuberculosis* during the non-replicative state. The crystal structure of FolP1 in complex with 6-hydroxymethylpterin monophosphate (92) shares four interacting residues with decoquinate RMB041 (**Error! Reference source not found.**).

## Conclusion

Various *in silico* drug discovery strategies have been implemented during this study to identify potential drug targets for decoquinate RMB041 against *M. tuberculosis*. Furthermore, we generate additional support for the previously determined metabolic pathways that were disrupted by decoquinate RMB041 and indicate at which enzymes the particular disruptions may occur. In addition, feasible synergistic activity of decoquinate RMB041 with other antimicrobials was shown. In this study, we also elaborated on the possible activity of decoquinate RMB041 against dormant *M. tuberculosis* and identified the drug targets for such. Lastly, the importance of including *in silico* drug discovery strategies and how they can be used as a complementary tool to other research approaches (*in vivo and in vitro* techniques) is highlighted.

## Methods

To ensure the probability of high effectivity and low toxicity of decoquinate RMB041, several tools were employed. The procedure followed during this study is illustrated in Figure 7.

**Figure 6.**
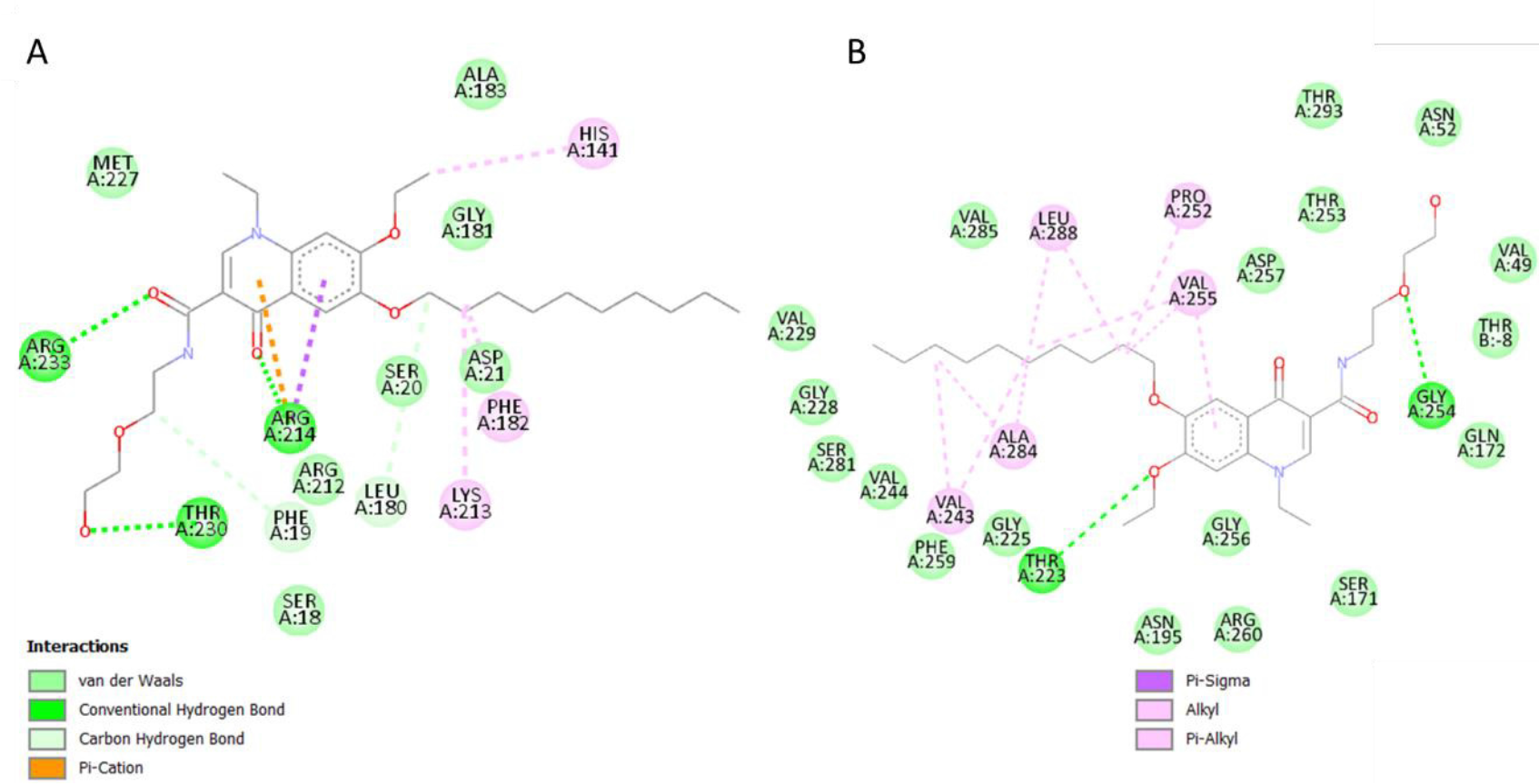
Interactions between decoquinate RMB041 and (A) AdoK and (B) FolP1.

**Figure 7.**
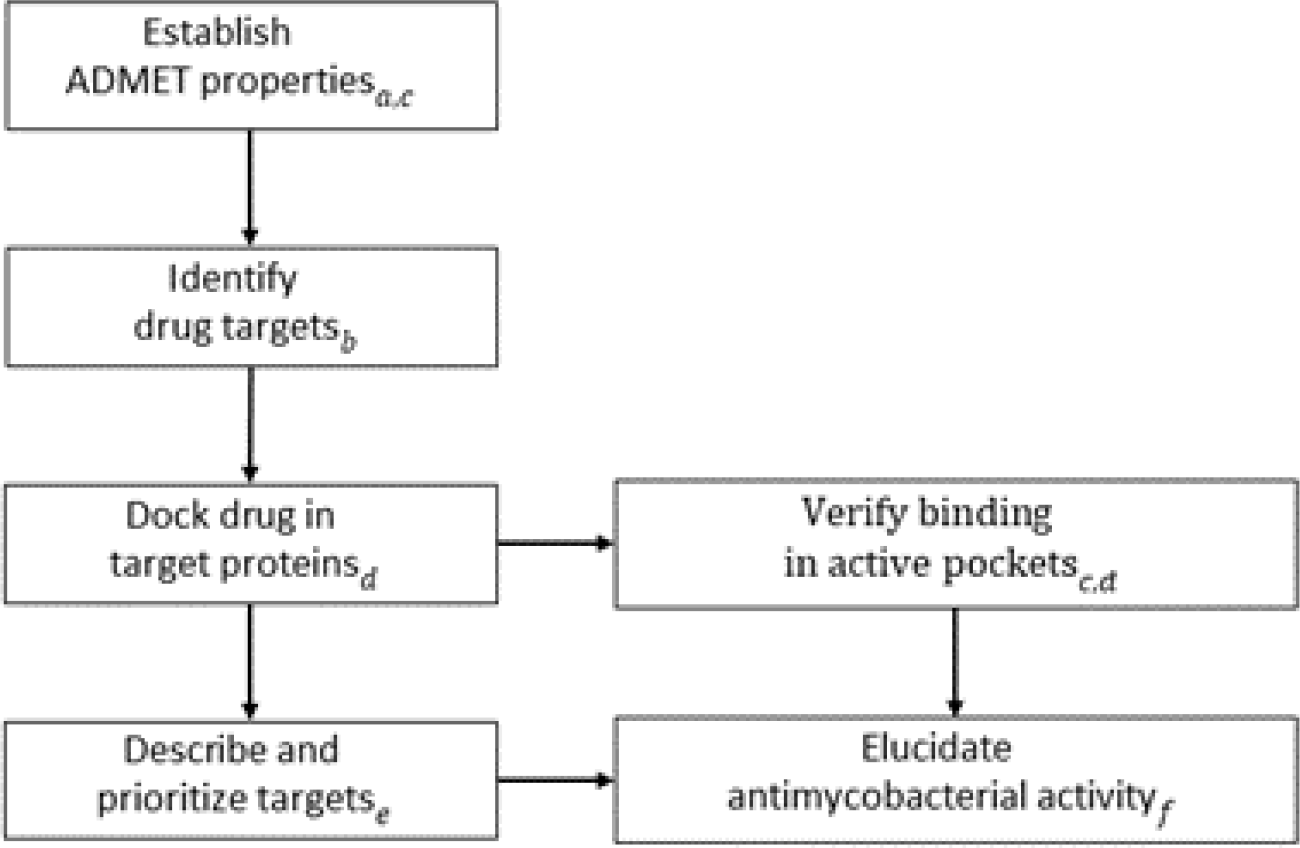
The workflow followed to elucidate the potential antimycobacterial mechanism of decoquinate derivative RMB041. The tools used during each step: a) pkCSM and SWISSadme, b) PharmMapper, c) Discovery Studio, d) AutoDock Vina, e) TDR Targets, f) KEGG and STRING.

### Evaluation of molecular and pharmakokinetic properties

The chemical 2D structure of the ligand was drawn with ACD/ChemSketch (commercial version) and converted to a 3D (pdb) file with DS v.4.5. ADMET properties of decoquinate RMB041 were identified by using the ADMET descriptors algorithm and toxicity prediction extensible protocol of BIOVIA DS Visualizer v.4.5 (Accelrys) (Software Inc., San Diego, CA) and the web servers Molinspiration (http://www.molinspiration.com//cgi-bin/properties), SwissADME (http://www.swissadme.ch/) and pkCSM (http://biosig.unimelb.edu.au/. The results are documented in **Error! Reference source not found.**. Although sharing similar functions, these tools apply different calculating algorithms, and can be used in combination to ensure improved precision of elucidation. SwissADME and DS also provided the molecular properties, which are reported in **Error! Reference source not found.**. The structure of decoquinate RMB041 was also evaluated with respect to pharmacokinetic rules frequently applied during drug manufacturing using this methodology (Table 2).

### Computational target fishing

#### Pharmacophore screening

PharmMapper (http://www.lilab-ecust.cn/pharmmapper/) was chosen in this study for the identification of potential protein targets, due to its wide popularity of use in applications of drug discovery. PharmMapper analyzes the spatial arrangement of key functional groups of a molecule and assigns pairwise fit scores according to the matching of pharmacophores between the ligand (being the drug) and the protein (14). For this study, protein targets with fitness scores of ≥4 were chosen to filter out insignificant pharmacophore models. The pharmacophores are collected in an internal database, PharmTargetDB, annotated from continuously updated databases, DrugBank, BindingDB, PDTD, TargetBank and Protein Database Bank (PDB). This software uses Cavity to identify all potential binding sites in proteins. The identification codes were retrieved from UniProt (https://www.uniprot.org/uniprot/) and the PDB (https://www.rcsb.org/) database. Only predicted protein targets relevant to *M. tuberculosis* were included for evaluation and deemed of importance of the study, and those in humans and other bacteria were excluded. The results are documented in **Table** .

### Reverse Docking

To validate the identified drug targets, decoquinate RMB041 was docked in the active cavity of each of PharmMapper’s predicted target proteins deemed valuable for the treatment of TB. The targets were then arranged according to their estimated binding strengths. Results documented in **Table** .

#### Protein and ligand preparation

PDB codes provided by PharmMapper were searched in PDB (http://www.pdb.org), where their three-dimensional structures with their co-crystalized ligands (solved at 0.90-1.8 Å resolution^2^) were then downloaded. Manual preparation was done with AutoDock v.4.2.6, which included the removal of water, deletion of all hetero atoms, assignment of atom types, addition of polar hydrogen atoms, merging of non-polar hydrogen atoms, and finally, the addition of Kollmann charges. Although docking programs account for flexibility of ligands, a remaining challenge is the flexibility of the entire protein. To minimize standard errors, the proteins were prepared as rigid structures (93). A pdbqt file of each of the target proteins were subsequently prepared with AutoDock Tools v.1.5.6 (94). Hydrogen and partial charge in the molecular system were assigned using AMBER force field.

For the preparation of decoquinate RMB041, its geometry force field was minimized using YASARA (http://www.yasara.org/) (95), prior to manual preparation involving the addition of polar hydrogens atoms, merging of non-polar hydrogens atoms, and addition of Gasteiger charges. PDBQT files were saved for docking.

#### Grid box preparation and docking

File conversions (mol2 to pdb to pdbqt) required for the separate steps of docking were performed with the open-source toolbox Open Babel v. 2.3.2 (96). To ensure inclusion of all active sites, the entire macromolecule was selected as a search space for binding site cavities, with spacing’s set at 0.90-1 Å, and saved as grid box parameters. Molecular docking scores of the compound to each protein were calculated by AutoDock Vina v.1.1.2, for which a Lamarckian Genetic Algorithm was used, and expressed as free binding energies (ΔG kcal·mol^-1^) (97). Nine different orientations of decoquinate RMB041 were searched per protein and ranked according to the binding free energies. To identify configurations that bind to active sites of their respective proteins, each binding pose was visually inspected in AutoDock, and those that interact with active site residues were saved as a pdbqt file and analyzed in DS, where the residues were then labeled. To validate interactions of decoquinate RMB041 within active cavity sites, the target residues were compared to those of the original co-crystalized ligands, which, if PDB cavity site records were available, were either retrieved with DS Tools v.1.5.6 or obtained from previous literature. Cavity sited were provided with the name of the specific amino acid, the respective chain, and the number of the residues, e.g., Lys A:123.

### Prioritization of protein targets

To identify proteins that are most likely essential during infection, i.e., important during the *M tuberculosis* transition to dormancy/latency, the structure of the compound was uploaded to the webserver TDR Targets (https://tdrtargets.org). TDR Targets, first introduced in 2008, has since been a reliable open-access resource for finding and prioritizing novel protein targets (46). This webserver integrates genomes from EupathDB, GenBank, GenoList and Mycobrowser and isolates proteins based on assay space and literature reviews on essential proteins during survival during dormancy/latency (e.g., by lack of oxygen, lack of nutrients, ROS, acidity).

### KEGG, GO, and network analysis

Annotations of the target proteins were retrieved from Kyoto Encyclopedia of Genes and Genomes (KEGG)(https://www.genome.jp/kegg/). A literature search of the identified target proteins of both active and dormant *M. tuberculosis* was done. Further, to include up to date information on dormant *M. tuberculosis*, articles published within the past 5 years were selected with the exclusion of preprints, articles citing original articles, and studies on vaccinations and diagnostics.

### Co-expression

Proteins that are negatively and positively correlated to any of the selected hit targets (Pearson correlation coefficient < -0.6 and > 0.6) (98), were obtained from TBDB (http://tbdb.bu.edu/). Proteins without known functions or assigned COG (Clusters of Orthologous Groups) categories were excluded. Target proteins were entered into STRING (https://string-db.org/), which provided a network with protein-protein interactions (medium confidence score of 0.4) in *M. tuberculosis* H37Rv.

## Acknowledgements

The authors greatly thank Jacques Petzer, Richard M. Beteck and Richard K. Haynes from the Department of Pharmaceutical Chemistry (NWU) for their assistance during the docking.

## References

1. WHO. Global tuberculosis report 2020: World Health Organization Geneva2020 [Available from: https://creativecommons.org/licenses/by-nc-sa/3.0/igo).

2. WHO. World Health Organizationon, Consolidated guidelines on drug-resistant tuberculosis treatment. Geneva. 2019.

3. Martinez-Jimenez F, Papadatos G, Yang L, Wallace IM, Kumar V, Pieper U, et al. Target prediction for an open access set of compounds active against Mycobacterium tuberculosis. PLoS Comput Biol. 2013;9(10):e1003253.

4. Mueller EA, Levin PA. Bacterial Cell Wall Quality Control during Environmental Stress. Molec Biol Physiol. 2020;11:e02456–20.

5. Stelitano G, Sammartino JC, Chiarelli LR. Multitargeting compounds: a promising strategy to overcome multi-drug resistant tuberculosis. Molecules. 2020;25(5):1239.

6. Andryukov B, Somova L, Matosova E, Lyapun I. Phenotypic plasticity as a strategy of bacterial resistance and an object of advanced antimicrobial technologies. Современные технологии в медицине. 2019;11(2).

7. Gample SP, Agrawal S, Sarkar D. Evidence of nitrite acting as a stable and robust inducer of non-cultivability in Mycobacterium tuberculosis with physiological relevance. Sci Rep. 2019;9(1):9261.

8. Campbell AJ, Lamb ML, Joseph-McCarthy D. Ensemble-based docking using biased molecular dynamics. J Chem Inf Model. 2014;54(7):2127–38.

9. Hurle M, Yang L, Xie Q, Rajpal D, Sanseau P, Agarwal P. Computational drug repositioning: from data to therapeutics. Clinical Pharmacology & Therapeutics. 2013;93(4):335–41.

10. Xu H, Jiao Y, Qin S, Zhao W, Chu Q, Wu K. Organoid technology in disease modelling, drug development, personalized treatment and regeneration medicine. Exp Hemat Oncol. 2018;7(1):1–12.

11. Knoll KE, Lindeque Z, Adeniji AA, Oosthuizen CB, Lall N, Loots DT. Elucidating the Antimycobacterial Mechanism of Action of Decoquinate Derivative RMB041 Using Metabolomics. Antibiotics. 2021;10(6):693.

12. Beteck RM, Seldon R, Coertzen D, van der Watt ME, Reader J, Mackenzie JS, et al. Accessible and distinct decoquinate derivatives active against Mycobacterium tuberculosis and apicomplexan parasites. Comm Chem. 2018;1(1).

13. Tanner L, Haynes RK, Wiesner L. An in vitro ADME and in vivo Pharmacokinetic Study of Novel TB-Active Decoquinate Derivatives. Front Pharmacol. 2019;10:120.

14. Wang X, Shen Y, Wang S, Li S, Zhang W, Liu X, et al. PharmMapper 2017 update: a web server for potential drug target identification with a comprehensive target pharmacophore database. Nucleic Acids Res. 2017;45(W1):W356–W60.

15. Vieira TF, Sousa SF. Comparing AutoDock and Vina in Ligand/Decoy Discrimination for Virtual Screening. Appl Sci. 2019;9(21).

16. Dar AM, Mir S. Molecular Docking: Approaches, Types, Applications and Basic Challenges. J Analyt Bioanalyt Tech. 2017;08(02).

17. Ali MT, Blicharska N, Shilpi JA, Seidel V. Investigation of the anti-TB potential of selected propolis constituents using a molecular docking approach. Sci Rep. 2018;8(1):12238.

18. Pagadala NS, Syed K, Tuszynski J. Software for molecular docking: a review. Biophys Rev. 2017;9(2):91–102.

19. Pantsar T, Poso A. Binding Affinity via Docking: Fact and Fiction. Molecules. 2018;23(8).

20. Macalino SJY, Billones JB, Organo VG, Carrillo MCO. In Silico Strategies in Tuberculosis Drug Discovery. Molecules. 2020;25(3).

21. Mishra H, Singh N, Lahiri T, Misra K. A comparative study on the molecular descriptors for predicting drug-likeness of small molecules. Bioinformation. 2009;3(9):384.

22. Mullard A. Re-assessing the rule of 5, two decades on. Nat Rev Drug Discov. 2018;17(11):777.

23. Lipinski CA. Drug-like properties and the causes of poor solubility and poor permeability. Journal of pharmacological and toxicological methods. 2000;44(1):235–49.

24. Veber DF, Johnson SR, Cheng H-Y, Smith BR, Ward KW, Kopple KD. Molecular properties that influence the oral bioavailability of drug candidates. J Med Chem. 2002;45(12):2615–23.

25. Egan WJ, Merz KM, Baldwin JJ. Prediction of drug absorption using multivariate statistics. J Med Chem. 2000;43(21):3867–77.

26. Muegge I, Heald SL, Brittelli D. Simple selection criteria for drug-like chemical matter. J Med Chem. 2001;44(12):1841–6.

27. Iacobino A, Piccaro G, Giannoni F, Mustazzolu A, Fattorini L. Fighting tuberculosis by drugs targeting nonreplicating Mycobacterium tuberculosis bacilli. Int J Mycobacteriol. 2017;6(3):213.

28. Tihanyi KK, Vastag M. Solubility, delivery and ADME problems of drugs and drug candidates: Bentham Science Publishers; 2011.

29. Bergström CA, Avdeef A. Perspectives in solubility measurement and interpretation. ADMET and DMPK. 2019;7(2):88–105.

30. Das MK, Chakraborty T. Progress in brain delivery of anti-HIV drugs. J Appl Pharm Sci. 2015;5(07):154–64.

31. Rana F. Rifampicin—an overview. Int J Res Pharm Chem. 2013;3(1):83–7.

32. Bangert MK, Hasbun R. Neurological and Psychiatric Adverse Effects of Antimicrobials. CNS Drugs. 2019;33(8):727–53.

33. Tanner L, Haynes RK, Wiesner L. Accumulation of TB-Active Compounds in Murine Organs Relevant to Infection by Mycobacterium tuberculosis. Front Pharmacol. 2020;11:724.

34. Smith DA, Beaumont K, Maurer TS, Di L. Volume of distribution in drug design: Miniperspective. J Med Chem. 2015;58(15):5691–8.

35. Glue P, Clement RP. Cytochrome P450 enzymes and drug metabolism—basic concepts and methods of assessment. Cell Mol Neurobiol. 1999;19(3):309–23.

36. Ji B, Truffot-Pernot C, Lacroix C, Raviglione MC, O’Brien RJ, Olliaro P, et al. Effectiveness of rifampin, rifabutin and rifapentine for preventive therapy of tuberculosis in mice. Am J Resp Critic Care Med. 1993;148(6):1541–6.

37. Jayaram R, Shandil RK, Gaonkar S, Kaur P, Suresh B, Mahesh B, et al. Isoniazid pharmacokinetics-pharmacodynamics in an aerosol infection model of tuberculosis. Antimicrob Agents Chemother. 2004;48(8):2951–7.

38. Arbex MA, Varella MdCL, Siqueira HRd, Mello FAFd. Antituberculosis drugs: drug interactions, adverse effects, and use in special situations-part 2: second line drugs. J Brasileiro de Pneumologia. 2010;36(5):641–56.

39. Ray A, Nangia V, Chatterji RS, Dalal N, Ray RS. Recurrent heart failure in pulmonary tuberculosis patients on antitubercular therapy: A case of protector turning predator. Egyp J Bronchol. 2017;11(3):288–91.

40. Kwon BS, Kim Y, Lee SH, Lim SY, Lee YJ, Park JS, et al. The high incidence of severe adverse events due to pyrazinamide in elderly patients with tuberculosis. PloS one. 2020;15(7):e0236109.

41. Lee N, Nguyen H. Ethambutol. StatPearls [Internet]. Treasure Island (FL): StatPearls Publishing; 2021 Jan.

42. Polianciuc SI, Gurzau AE, Kiss B, Stefan MG, Loghin F. Antibiotics in the environment: causes and consequences. Med Pharm Rep. 2020;93(3):231–40.

43. Hopkins AL. Network pharmacology. Nature biotechnology. 2007;25(10):1110–1.

44. Anand P, Chandra N. Characterizing the pocketome of Mycobacterium tuberculosis and application in rationalizing polypharmacological target selection. Scientific reports. 2014;4(1):1–17.

45. Melak T, Gakkhar S. Comparative genome and network centrality analysis to identify drug targets of Mycobacterium tuberculosis H37Rv. BioMed Res Int. 2015;2015:212061.

46. Landaburu L, Berenstein AJ, Videla S, Maru P, Shanmugam D, Chernomoretz A, et al. TDR Targets 6: driving drug discovery for human pathogens through intensive chemogenomic data integration. Nucleic Acids Res. 2020;48(D1):D992–D1005.

47. Hasan S, Daugelat S, Rao PSS, Schreiber M. Prioritizing Genomic Drug Targets in Pathogens: Application to Mycobacterium tuberculosis. PLoS Comp Biol. 2006;2(6).

48. Murphy DJ, Brown JR. Identification of gene targets against dormant phase Mycobacterium tuberculosis infections. BMC Infect Dis. 2007;7:84.

49. Papakonstantinou D, Dunn SJ, Draper SJ, Cunningham AF, O’Shea MK, McNally A. Mapping Gene-by-Gene Single-Nucleotide Variation in 8,535 Mycobacterium tuberculosis Genomes: a Resource To Support Potential Vaccine and Drug Development. mSphere. 2021;6(2).

50. Galagan JE, Sisk P, Stolte C, Weiner B, Koehrsen M, Wymore F, et al. TB database 2010: overview and update. Tuberculosis. 2010;90(4):225–35.

51. Mishra S, Shukla P, Bhaskar A, Anand K, Baloni P, Jha RK, et al. Efficacy of beta-lactam/beta-lactamase inhibitor combination is linked to WhiB4-mediated changes in redox physiology of Mycobacterium tuberculosis. Elife. 2017;6.

52. Sala C, Haouz A, Saul FA, Miras I, Rosenkrands I, Alzari PM, et al. Genome-wide regulon and crystal structure of BlaI (Rv1846c) from Mycobacterium tuberculosis. Mol Microbiol. 2009;71(5):1102–16.

53. Black PA, Warren RM, Louw GE, van Helden PD, Victor TC, Kana BD. Energy metabolism and drug efflux in Mycobacterium tuberculosis. Antimicrob Agents Chemother. 2014;58(5):2491–503.

54. Podust LM, Ouellet H, von Kries JP, de Montellano PRO. Interaction of Mycobacterium tuberculosis CYP130 with heterocyclic arylamines. J Biol Chem. 2009;284(37):25211–9.

55. Ouellet H, Podust LM, de Montellano PR. Mycobacterium tuberculosis CYP130: crystal structure, biophysical characterization, and interactions with antifungal azole drugs. J Biol Chem. 2008;283(8):5069–80.

56. Bacon J, Alderwick LJ, Allnutt JA, Gabasova E, Watson R, Hatch KA, et al. Non-replicating Mycobacterium tuberculosis elicits a reduced infectivity profile with corresponding modifications to the cell wall and extracellular matrix. PLoS One. 2014;9(2):e87329.

57. Argyrou A, Vetting MW, Blanchard JS. Characterization of a new member of the flavoprotein disulfide reductase family of enzymes from Mycobacterium tuberculosis. J Biol Chem. 2004;279(50):52694–702.

58. Zhai W, Wu F, Zhang Y, Fu Y, Liu Z. The immune escape mechanisms of Mycobacterium tuberculosis. Int J Molec Sci. 2019;20(2):340.

59. Sharma V, Arockiasamy A, Ronning DR, Savva CG, Holzenburg A, Braunstein M, et al. Crystal structure of Mycobacterium tuberculosis SecA, a preprotein translocating ATPase. PNAS. 2003;100:2243–8.

60. Duong F. Binding, activation and dissociation of the dimeric SecA ATPase at the dimeric SecYEG translocase. The EMBO Journal. 2003;22(17):4375–84.

61. Hou JM, D’lima NG, Rigel NW, Gibbons HS, McCann JR, Braunstein M, et al. ATPase activity of Mycobacterium tuberculosis SecA1 and SecA2 proteins and its importance for SecA2 function in macrophages. J Bacteriol. 2008;190(14):4880–7.

62. Muller AU, Weber-Ban E. The Bacterial Proteasome at the Core of Diverse Degradation Pathways. Front Mol Biosci. 2019;6:23.

63. Raman K, Chandra N. Mycobacterium tuberculosis interactome analysis unravels potential pathways to drug resistance. BMC Microbiol. 2008;8:234.

64. Schwab U, Rohde KH, Wang Z, Chess PR, Notter RH, Russell DG. Transcriptional responses of Mycobacterium tuberculosis to lung surfactant. Microbial pathogenesis. 2009;46(4):185–93.

65. Sulzenbacher G, Canaan S, Bordat Y, Neyrolles O, Stadthagen G, Roig-Zamboni V, et al. LppX is a lipoprotein required for the translocation of phthiocerol dimycocerosates to the surface of Mycobacterium tuberculosis. EMBO J. 2006;25(7):1436–44.

66. Nicoara SC, Minnikin DE, Lee OC, O’Sullivan DM, McNerney R, Pillinger CT, et al. Development and optimization of a gas chromatography/mass spectrometry method for the analysis of thermochemolytic degradation products of phthiocerol dimycocerosate waxes found in Mycobacterium tuberculosis. Rapid Commun Mass Spectrom. 2013;27(21):2374–82.

67. Crellin PK, Luo C-Y, Morita YS. Metabolism of Plasma Membrane Lipids in Mycobacteria and Corynebacteria. Lipid Metabolism. 2013.

68. Slayden RA, Barry CE, 3rd. The role of KasA and KasB in the biosynthesis of meromycolic acids and isoniazid resistance in Mycobacterium tuberculosis. Tuberculosis (Edinb). 2002;82(4-5):149–60.

69. Meneguello JE, Arita GS, de Oliveira Silva JV, Ghiraldi-Lopes LD, Caleffi-Ferracioli KR, Siqueira VLD, et al. Insight about cell wall remodulation triggered by rifampicin in Mycobacterium tuberculosis. Tuberculosis. 2020;120:101903.

70. Jagadeb M, Rath SN, Sonawane A. In silico discovery of potential drug molecules to improve the treatment of isoniazid-resistant Mycobacterium tuberculosis. Journal of Biomolecular Structure and Dynamics. 2019;37(13):3388–98.

71. Nieto LM, Mehaffy C, Dobos KM. The physiology of mycobacterium tuberculosis in the context of drug resistance: a system biology perspective. Mycobacterium-Research and Development. 2017.

72. Kumari R, Saxena R, Tiwari S, Tripathi DK, Srivastava KK. Rv3080c regulates the rate of inhibition of mycobacteria by isoniazid through FabD. Mol Cell Biochem. 2013;374(1):149–55.

73. Festa RA, McAllister F, Pearce MJ, Mintseris J, Burns KE, Gygi SP, et al. Prokayrotic ubiquitin-like protein (Pup) proteome of Mycobacterium tuberculosis. PloS one. 2010;5(1):e8589.

74. Chakraborty P, Kumar A. The extracellular matrix of mycobacterial biofilms: could we shorten the treatment of mycobacterial infections? Microb Cell. 2019;6(2):105–22.

75. Li Z, Huang Y, Ge J, Fan H, Zhou X, Li S, et al. The crystal structure of MCAT from Mycobacterium tuberculosis reveals three new catalytic models. J Mol Biol. 2007;371(4):1075–83.

76. Trutneva KA, Shleeva MO, Demina GR, Vostroknutova GN, Kaprelyans AS. One-Year Old Dormant, “Non-culturable” Mycobacterium tuberculosis Preserves Significantly Diverse Protein Profile. Front Cell Infect Microbiol. 2020;10:26.

77. Wilkinson RJ, DesJardin LE, Islam N, Gibson BM, Kanost RA, Wilkinson KA, et al. An increase in expression of a Mycobacterium tuberculosis mycolyl transferase gene (fbpB) occurs early after infection of human monocytes. Mol Microbiol. 2001;39(3):813–21.

78. Anderson DH, Harth G, Horwitz MA, Eisenberg D. An interfacial mechanism and a class of inhibitors inferred from two crystal structures of the Mycobacterium tuberculosis 30 kDa major secretory protein (Antigen 85B), a mycolyl transferase. J Mol Biol. 2001;307(2):671–81.

79. Betts JC, McAdam RA, Lukey PT, Duncan K. Evaluation of a nutrient starvation model of Mycobacterium tuberculosis persistence by gene and protein expression profiling. Molec Microbiol. 2002;43:717–31.

80. Huang H-L, Krieger IV, Parai MK, Gawandi VB, Sacchettini JC. Mycobacterium tuberculosis malate synthase structures with fragments reveal a portal for substrate/product exchange. J Biol Chem. 2016;291(53):27421–32.

81. Hamidieh F, Farnia P, Nowroozi J, Farnia P, Velayati AA. An Overview of Genetic Information of Latent Mycobacterium tuberculosis. TB Resp Dis. 2021;84(1):1.

82. Smith CV, Huang CC, Miczak A, Russell DG, Sacchettini JC, Honer zu Bentrup K. Biochemical and structural studies of malate synthase from Mycobacterium tuberculosis. J Biol Chem. 2003;278(3):1735–43.

83. Rodionov DA, Vitreschak AG, Mironov AA, Gelfand MS. Regulation of lysine biosynthesis and transport genes in bacteria: yet another RNA riboswitch? Nucl Acids Res. 2003;31(23):6748–57.

84. Gokulan K, Rupp B, Pavelka MS, Jr., Jacobs WR, Jr., Sacchettini JC. Crystal structure of Mycobacterium tuberculosis diaminopimelate decarboxylase, an essential enzyme in bacterial lysine biosynthesis. J Biol Chem. 2003;278(20):18588–96.

85. Moreira C, Ramos MJ, Fernandes PA. Glutamine synthetase drugability beyond its active site: exploring oligomerization interfaces and pockets. Molecules. 2016;21(8):1028.

86. Krajewski WW, Jones AT, Mowbray SL. Structure of Mycobacterium tuberculosis glutamine synthetase in complex with a transition-state mimic provides functional insights. PNAS. 2005;102:10499–504.

87. Agapova A, Serafini A, Petridis M, Hunt DM, Garza-Garcia A, Sohaskey CD, et al. Flexible nitrogen utilisation by the metabolic generalist pathogen Mycobacterium tuberculosis. Elife. 2019;8.

88. Tullius MV, Harth G, Horwitz MA. Glutamine synthetase GlnA1 is essential for growth of Mycobacterium tuberculosis in human THP-1 macrophages and guinea pigs. Infect Immun. 2003;71(7):3927–36.

89. Long MC, Escuyer V, Parker WB. Identification and characterization of a unique adenosine kinase from Mycobacterium tuberculosis. Journal of bacteriology. 2003;185(22):6548–55.

90. Reddy MCM, Palaninathan SK, Shetty ND, Owen JL, Watson MD, Sacchettini JC. High resolution crystal structures of Mycobacterium tuberculosis adenosine kinase: insights into the mechanism and specificity of this novel prokaryotic enzyme. J Biol Chem. 2007;282(37):27334–42.

91. Liu T, Wang B, Guo J, Zhou Y, Julius M, Njire M, et al. Role of folP1 and folP2 genes in the action of sulfamethoxazole and trimethoprim against mycobacteria. J Microbiol Biotech. 2015;25(9):1559–67.

92. Baca AM, Sirawaraporn R, Turley S, Sirawaraporn W, Hol WG. Crystal structure of Mycobacterium tuberculosis 7,8-dihydropteroate synthase in complex with pterin monophosphate: new insight into the enzymatic mechanism and sulfa-drug action. J Mol Biol. 2000;302(5):1193–212.

93. Trott O, Olson AJ. AutoDock Vina: improving the speed and accuracy of docking with a new scoring function, efficient optimization, and multithreading. J Comp Chem. 2010;31(2):455–61.

94. Huey R, Morris GM, Forli S. Using AutoDock 4 and AutoDock vina with AutoDockTools: a tutorial. The Scripps Research Institute Mol Graph Lab. 2012;10550:92037.

95. Land H, Humble MS. YASARA: a tool to obtain structural guidance in biocatalytic investigations. Protein Engineering: Springer; 2018. p. 43–67.

96. O’Boyle NM, Banck M, James CA, Morley C, Vandermeersch T, Hutchison GR. Open Babel: An open chemical toolbox. J Cheminform. 2011;3(1):1–14.

97. Sandeep G, Nagasree KP, Hanisha M, Kumar MMK. AUDocker LE: A GUI for virtual screening with AUTODOCK Vina. BMC research notes. 2011;4(1):1–4.

98. Akoglu H. User’s guide to correlation coefficients. Turk J Emerg Med. 2018;18(3):91–3.

